# Paleocene origin of the Neotropical lineage of cleptoparasitc bees Ericrocidini-Rhathymini (Hymenoptera, Apidae)

**DOI:** 10.1101/224683

**Authors:** Aline C. Martins, David R. Luz, Gabriel A. R. Melo

**Author notes:** CORRESPONDING AUTHOR: Aline C. Martins EMAIL OF CORRESPONDING AUTHOR.

## Abstract

Cleptoparasitic bees abandoned the pollen collecting for their offspring and lay their eggs on other bees’ provisioned nests. Also known as cuckoo bees they belong to several lineages, especially diverse in Apinae. We focused on a lineage of Apinae cleptoparasitic bees, the clade Ericrocidini+Rhathymini, which attack nests of the oil-collecting bees. We sequenced five genes for a broad sampling in this clade plus a large outgroup and reconstruct phylogeny and divergence times. We confirmed the monophyly of the clade Ericrocidini+Rhathymini and its position inside the ericrocidine line, together with the tribes Protepeolini, Isepeolini and Coelioxoidini. Our results corroborate the current taxonomic classification. *Ericrocis* is the basal most lineage in Ericrocidini and the position of *Acanthopus* and the most diverse genus *Mesoplia* were inconclusive. Ericrocidini+Rhathymini diverged from *Parepeolus aterrimus* 74 mya in the Cretaceous. Considering the robust molecular evidence of their sister relationships, the striking differences on the first instar larvae morphology of the two groups are probably adaptations to the distinct nesting biology of their hosts. As other parasites in the ericrocidine line, both groups possess larvae adapted to kill the immature host and to feed on floral oil provisioned by the host female. The evolution of host specialization in the line Ericrocidini+Rhathymini retroced to the Eocene when they arose synchronously with their hosts, *Centris* and *Epicharis*.

## Introduction

Bees are the most important lineage of pollinating animals acting directly in the reproductive success of wild and cultivated angiosperms, therefore playing an important role in environment and human food production (Kremen et al., 2002; Klein et al., 2007). The origin of the bee lineage in the early Cretaceous, approximately 120 million years ago, is coincident with radiation of the most representative group of angiosperms, the eudicots, which are heavily dependent on bees for pollination (Cardinal & Danforth, 2013). The feeding behavior of bees represents a novelty in the evolutionary history of Hymenoptera: the changing from an carnivorous diet present in bee’s closest relatives—apoid wasps and ants—to an exclusively vegetarian diet based on pollen feeding (Melo et al., 2011; Peters et al., 2017). Female and male bee adults feed on nectar produced by flowers, while the young are feed mainly with pollen as a source of protein, together with other floral energy-rich resources like nectar and oil (Michener, 2007).

The almost 20,000 bee species are recognized as belonging to a single family, the Apidae s.l. (Melo & Goncalves, 2005), classified in seven lineages—Andreninae, Apinae, Colletinae, Megachilinae, Melittinae and Stenotritinae—and distributed in all continents except Antarctica (Michener, 2007). The monophyly of Apidae is undoubtful and corroborated by morphological, behavioral and molecular evidences (Melo, 1999; Danforth et al., 2006; Michener, 2007; Branstetter et al., 2017; Peters et al., 2017).

Much progress has been made on our understanding of bee phylogenetic relationships by the increasing use of molecular data and model-based methods. As a consequence, the newly proposed bee phylogenies have shed light on our understanding of crucial aspects of bee evolution like plant host choice evolution, social behaviour and cleptoparasitism (Danforth et al., 2011).

All bees feed on flower resources, but females of many species abandoned both pollen collecting and construction of nests for their own offspring. Instead they parasitize other bee species laying their eggs in provisioned nests (Michener, 2007).

This relationship where the offspring of one species feed and develops in the food stored by a female of other species is called brood parasitism, or specifically in bees, cleptoparasitism or cleptoparasitic behavior (Rozen, 1991). Cleptoparasitism has been reported in almost all bee lineages, excepting Mellitinae, but a special diversity is found in the long-tongued lineage (subfamilies Apinae and Megachilinae) (Straka & Bogusch, 2007a; Cardinal et al., 2010; Litman et al., 2013). The cleptoparasitism originated early in the history of bees, approximately 95 Mya in the Cretaceous, in the large *cleptoclade* of Apinae (Cardinal et al., 2010). While many parasitic groups evolved from pollen-collecting ancestors, the way back—cleptoparasite evolving to pollen-collecting—was never reported (Litman et al., 2013). Despite the risks associated to obligatory parasitism and apparent decrease in diversification rates associated to these lineages, it evolved many times independently in bees (Michener, 2007; Litman et al., 2013)

Close phylogenetic proximity between hosts and cleptoparasites has been argued by previous authors (Wcislo & Cane, 1996; Michener, 2007), following a pattern known mostly as Emery’s rule in ants and wasps (Höldobler & Wilson 1990). After the work of Emery (1909), who based his conclusion on a limited number of ant species, LeMasne (1956) extrapolated this pattern to other Hymenoptera and named it as Emery’s rule. In bees we can find some groups of cleptoparasites closely related to their hosts, especially on younger cleptoparasitic clades (Litman et al., 2013), for example: *Aglae* and *Exaerete* on *Eufriesea* and *Eulaema* (Euglossini) (Cardinal et al., 2010); *Ctenoplectrina* on *Ctenoplectra* (Ctenoplectrini) (Schaefer & Renner, 2008). However, most cleptoparasitic groups, at least in Apinae s.l., contradict Emery’s rule. Instead, cleptoparasitism arose four times independently in this bee clade and most cleptoparasite and hosts lineages were shown not be closely related (Cardinal et al., 2010). Most apine cleptoparasites are included in a large monophyletic group, including the nomadine tribes (here referred as the nomadine line) and additional tribes previously spread apart within Apinae (*e.g*. Melectini, Osirini, Rhathymini, Ericrocidini, Isepeolini, Protepeolini), with the other three independent origins referring to the genera *Exaerete* and *Aglae*, in Euglossini, and *Ctenoplectrina*, in Ctenoplectrini. This arrangement contradicts previous morphological analyses based on adult and larval characters (Roig-Alsina & Michener, 1993; Straka & Bogusch, 2007a).

Morphological data also suggested a close relatedness between the cleptoparasitic Ericrocidini and Rhathymini and their hosts, *Centris* and *Epicharis* (traditionally placed in a single tribe, Centridini, but see Martins & Melo 2016), evidenced by many supposedly shared characters (Snelling & Brooks, 1985). Further phylogenetic analyses based on morphology (Roig-Alsina & Michener, 1993; Straka & Bogusch, 2007a) and molecules (Cardinal et al., 2010) did not support their close relationship. Alternatively morphological phylogenies have been pointed to sister relationship between Ericrocidini and Melectini (Roig-Alsina & Michener, 1993) or Ericrocidini and Isepeolini (Straka & Bogusch, 2007a). All recent molecular evidences contradict both hypothesis and pointed to the sister relationship between Ericrocidini and Rhathymini, which are nested inside the large cleptoclade, forming the ericrocidine line together with Protepeolini, Isepeolini and Osirini (Cardinal et al., 2010; Cardinal & Danforth, 2013; Litman et al., 2013), while *Centris* and *Epicharis* are closely related to the corbiculate bees (Martins et al., 2014; Martins & Melo, 2016).

Ericrocidini comprises approximately 44 species distributed in eleven genera, exhibiting mostly a neotropical distribution, with a single genus (*Ericrocis*) entering the Nearctic region, while Rhathymini includes only one genus, *Rhathymus*, with about 20 species, all neotropical (see Fig. 1 and 2 for morphological diversity in the group) (Moure & Melo, 2012a, 2012b). Snelling & Brooks (1985) analyzed the phylogenetic relationships among Ericrocidini genera, using a *bauplan* approach, with Rhathymini and Centridini as outgroups. Their phylogeny indicated Ericrocidini and Rhathymini as sister groups, although their outgroup choice was rather limited, and *Ericrocis* as the basal most lineage within Ericrocidini. Ericrocidini parasitize primarily nests of *Centris* (although *Mesoplia rufipes* can also attack nests of *Epicharis*; see Rocha-Filho et al., 2009 and references therein) and Rhathymini parasitize only species of *Epicharis* (Michener, 2007; Werneck et al., 2012).

**Figure 1.**
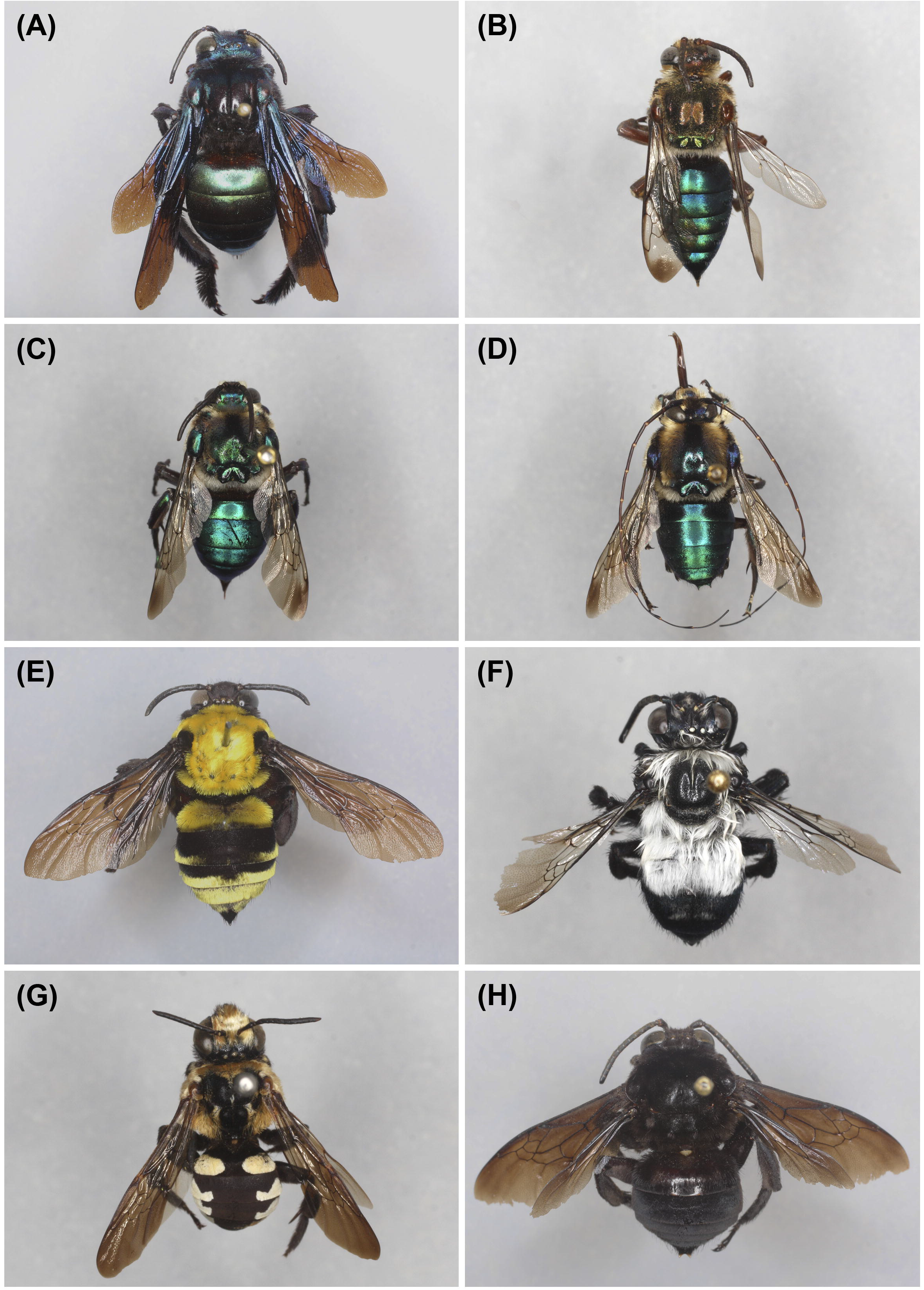
Diversity of Ericrocidini: a. *Acanthopus palmatus* (Olivier, 1789); b. *Aglaomelissa duckei* (Friese, 1906); c. *Ctenioschelus goryi* (Romand, 1840), female; d. *Ctenioschelus goryi*, male; e. *Cyphomelissa magnífica* Moure, 1958; f. *Epiclopus gayi* Spinola, 1851; g. *Ericrocis pintada* Snelling & Zavortink, 1985; h. *Eurytis funereus* Smith, 1854

**Figure 2.**
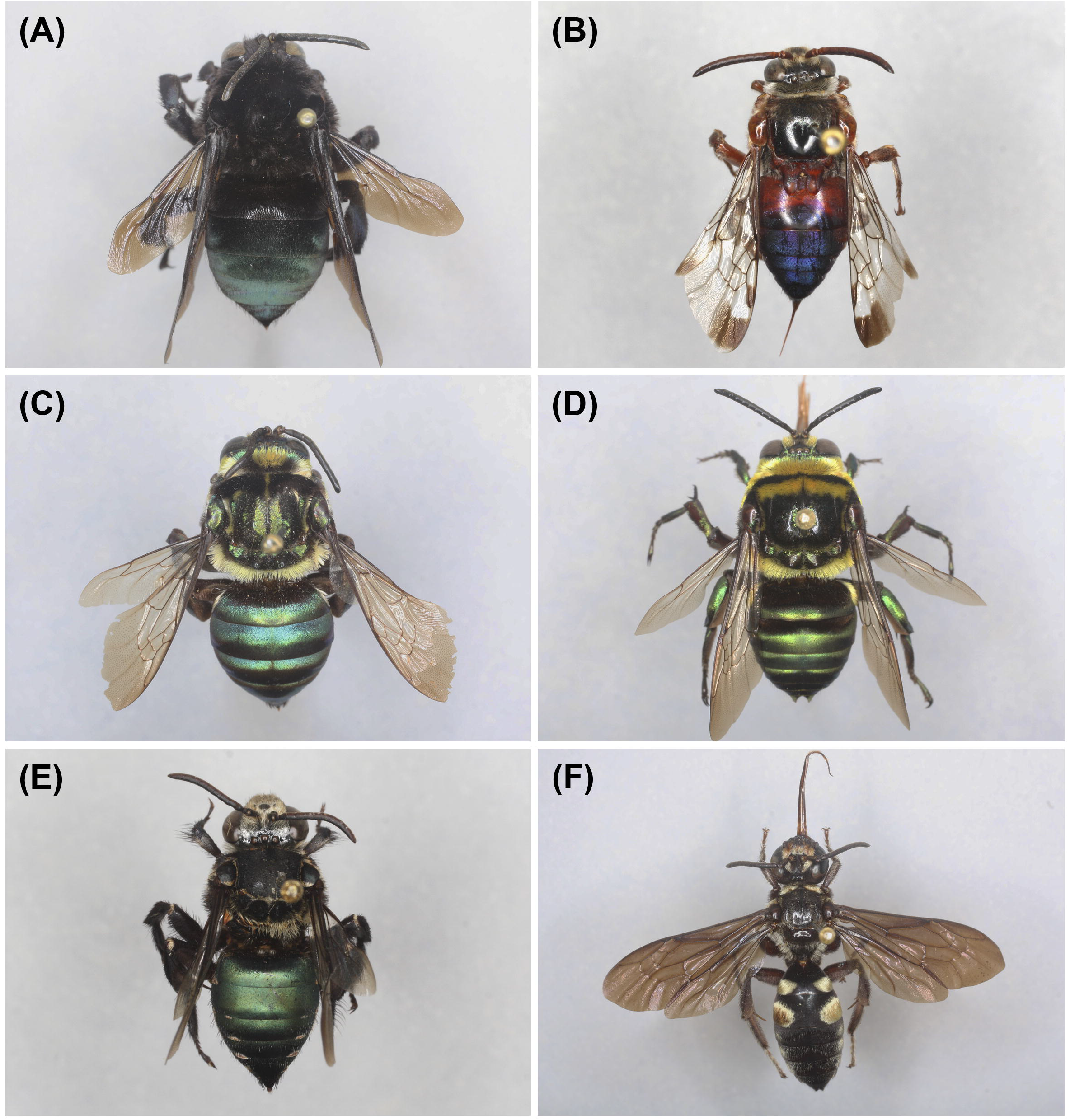
Diversity of Ericrocidini (a-e) and Rhathymini (f) genera: a. *Hopliphora velutina* Lepeletier & Serville, 1825; b. *Mesocheira bicolor* (Fabricius, 1804); c. *Mesoplia ornata* (Spinola, 1841), female; d. *Mesoplia ornata*, male; e. *Mesonychium coerulescens* Lepeletier & Serville, 1825; f. *Rhathymus quadriplagiatus* (Smith, 1860).

Host associations in Ericrocidini and Rhathymini are still poorly documented. The hosts of *Ctenioschelus, Eurytis* and *Hopliphora* remain unknown (Thiele, 2008; Rocha-Filho et al., 2009), and only recently it has been discovered that *Cyphomelissa* parasitizes *Centris (Melacentris)* (Rocha-Filho et al. 2017). Adult females of both tribes introduce their eggs into closed cells of the host by breaking a hole in the cell closure (Rozen, 2003; Rozen et al., 2011). When a female bee parasitizes a closed nest cell, her offspring will find the nest already provisioned and the young of the host female (egg or larva), which means the parasite larva needs to kill the host immature in order to have enough food for its development. This strategy involves morphological and behavioural adaptations on the cleptoparasitic larva (for example elongate, sickle-shaped mandibles) that allow them to kill the host offspring, therefore it is called hospicidal larva (from the latin *hospes=host, caedo=* to kill) (Rozen, 1989).

The present study focused on the phylogenetic relationships and divergence times among the clade Ericrocidini+Rhathymini, and the implications on classification and evolution of cleptoparasitic behaviour in this lineage. Taking into account recent hypotheses of relationship between Ericrocidini and Rhathymini, as well as their placement within Apinae, we reconstruct their phylogenetic relationships using related cleptoparasitic tribes from the ericrocidine and nomadine lines plus Melectini and Anthophorini as outgroups. This new outgroup choice associated to a more representative sampling will offer new opportunities to understanding the evolution of this bee group and to propose a phylogenetic based classification for Ericrocidini and Rhathymini.

## Material and Methods

### Taxon Sampling

We newly sequenced 17 species of Ericrocidini representing all genera, except *Aglaomelissa*, mostly with >2 species per genera, and five species of Rhathymini, totalizing 20 species and 10 genera (Table S1). We made an effort to include morphologically distinct species and a wider geographic distribution. In total, 73 new sequences have been submitted to GenBank (Table S1). Most of our ingroup representatives were newly sequenced, but we also included eight ingroup terminals from GenBank (Table S2). Vouchers from the newly produced sequences are deposited mostly at the DZUP – Jesus Santiago Moure Entomological Collection at Federal University of Paraná, Brazil, or at institutions that lent specimens from DNA extractions (Table S1). As outgroup, we included representatives of several tribes of Apinae (Anthophorini, Caenoprosopidini, Coelioxoidini, Epeolini, Ammobatoidini, Isepeolini, Protepeolini, Melectini, Nomadini, Osirini, Ammobatini), mostly cleoptoparasites from the cleptoclade plus Anthophorini, all downloaded from GenBank (Table S2). Taxonomic classification follows Moure et al. (2012).

### Molecular data sampling

DNA was isolated mostly from newly collected specimens preserved in EtOH, but also from some dried pinned specimens collected for less than 10 years. DNA was extracted using the Qiagen DNeasy blood & tissue extraction kit, following the manufacturer’s protocol, except for pinned specimens, which remained longer period in the lysis phase. We amplified and sequenced four nuclear genes: the ribosomal 28S gene (~1400 base pairs) and three nuclear protein-coding genes: LW-Rhodopsin (~700 base pairs), Elongation factor 1α – F2 copy (~1,000 base pairs), and RNA-polymerase (~900 base pairs). Additionally, we sequenced the mitochondrial barcode gene cytochrome oxidase I (~700 base pairs). PCR amplifications were performed with standard protocols. We used primers from the literature, but also designed two new primer pairs for LW-Rhodopsin and Elongation factor 1α to increase the success of amplification. All primers and PCR conditions are described in Table S3. PCR products were purified and sequenced by Macrogen Inc., South Korea. Forward and reverse strands were assembled, edited and analyzed in Geneious R8 (Biomatters, 2013). All sequences were Blast searched to prevent using contaminated samples. Sequences were submitted to GenBank using the GenBank submission tool plugin in Geneious R8.

### Alignment and phylogenetic inference

We aligned the sequences using the MAFFT vs. 7program (Katoh & Standley, 2013) using the parameters: 1PAM/k=2 for the nucleotide scoring matrix; 1.53 for gap opening penalty and 0 offset value (default); and leaving gappy regions. We used different alignment strategies depending on the gene. The 28S ribosomal gene were aligned using secondary structure and the algorithm Q-INS-i. The protein-coding genes that contain introns, *i.e*. elongation factor 1α and LW-Rhodopsin, were aligned using the E-INS-I strategy, recommended by MAFFT manual for sequences with multiple conserved domains and long gaps. The protein coding genes with no introns in the region we amplified (RNA polymerase) and the CO1 were aligned using G-INS-i, recommended for sequences with global homology. Except for the 28S structural alignment, which were performed on the MAFFT *online* server, all the remaining were performed in Geneious R8. Minor adjustments were made by eye. Each gene matrix was submitted to individual tree searches to check for strong incongruences among datasets (*i.e*. well-supported incongruences). In the absence of strongly supported incongruences all genes were concatenated in one matrix totalizing 4519 aligned nucleotides.

Dataset was partitioned in 13 partitions: the 28 S ribosomal gene; introns and exons of LW-Rhodopsin and elongation factor 1-alpha were considered as separated partitions, totalizing 8 partitions; RNA polymerase gene as one partition and the mitochondrial gene CO1 was partitioned by codon position. This scheme was used in the phylogenetic and dating analyses, except as indicated below.

Maximum likelihood tree searches and bootstrapping were performed in RaxML (Stamatakis, 2006) using the graphical interface raxmlGUI (Silvestro & Michalak, 2012) with 1000 bootstrap replicates. Bayesian tree searches were performed in MrBayes 3.2 (Ronquist et al., 2012) with 12 data partitions (the 3^rd^ position of the CO1 gene was excluded due to non-convergence caused by conflict with the other partitions during preliminary analyses). The Markov Chain Monte Carlo (MCMC) was run in the CIPRES server (Miller et al., 2010),for10 million generations, with trees sampled every 1000 generations. Default MrBayes priors were used, except for implementation of a mixed evolutionary model of nucleotide substitution. Convergence of the chains was assessed in Tracer 1.6 (Rambaut et al., 2014) and trees obtained prior to convergence were discarded as *burnin* (25%), and a 50% majority rule consensus tree was constructed from the remaining trees.

### Divergence times estimates

Bayesian age estimates were performed using BEAST 2 (Bouckaert et al., 2014) at the same matrix analyzed phylogenetically with Bayesian inference and maximum likelihood. We used a Yule tree prior, GTR+G substitution model and uncorrelated lognormal relaxed clock. The same partitioning scheme was applied, totalizing 13 partitions. The MCMC run was performed in the CIPRES server (Miller et al., 2010), with 50 million generations, sampled every 10000^th^ generations. Convergence of chains were checked in Tracer(BEAST package) considering effective sample sizes (ESS) values 200. The maximum clade credibility (MCC) tree was produced in TreeAnotator (BEAST package).

One fossil was used to calibrate the tree, *Paleohabropoda oudardi* from the Paleocene of Menat (Puy-de-Dôme, France), which is considered the third oldest bee fossil (Michez et al., 2009). Phylogenetic analysis of 17 morphological characters plus morphometric analysis indicated that this fossil clearly belongs to the extant tribe Anthophorini (Michez et al., 2009). *Paleohaproboda oudardi* have been used to calibrate the node uniting all Anthophorini (Cardinal et al., 2010; Cardinal & Danforth, 2013). Because it might represent the stem lineage of this tribe we applied a normal distribution prior with a mean of 60 Ma and stdev of 5. We also constrained the root age according to previous estimates for the divergence between Anthophorini and the cleptoclade (Martins & Melo, 2016) applying a normal distribution prior with mean of 100 Ma and stdev of 5. Fossil calibration and constrains are depicted in Figure 4.

## Results

### Phylogenetics

The aligned data matrix comprised 4419 nucleotides, of which 1283 were derived from the ribosomal gene 28S, 836 from RNA polymerase, 759 from long-wavelength rhodopsin, 963 from elongation-factor 1-alpha and 678 from the mitochondrial cytochrome oxidase I. Figure 3 and Supplementary Material (Figure S1) show the phylograms, respectively, of the 50% majority-rule consensus tree of the Bayesian analysis and the maximum likelihood tree. Both trees were congruent, *i.e* there were no clades that were strongly supported (≥70% bootstrap support, ≥0.95 Bayesian posterior probability) in one tree but contradicted in the other. Trees were deposited in TreeBase under number (will be submitted upon manuscript acceptance).

**Figure 3.**
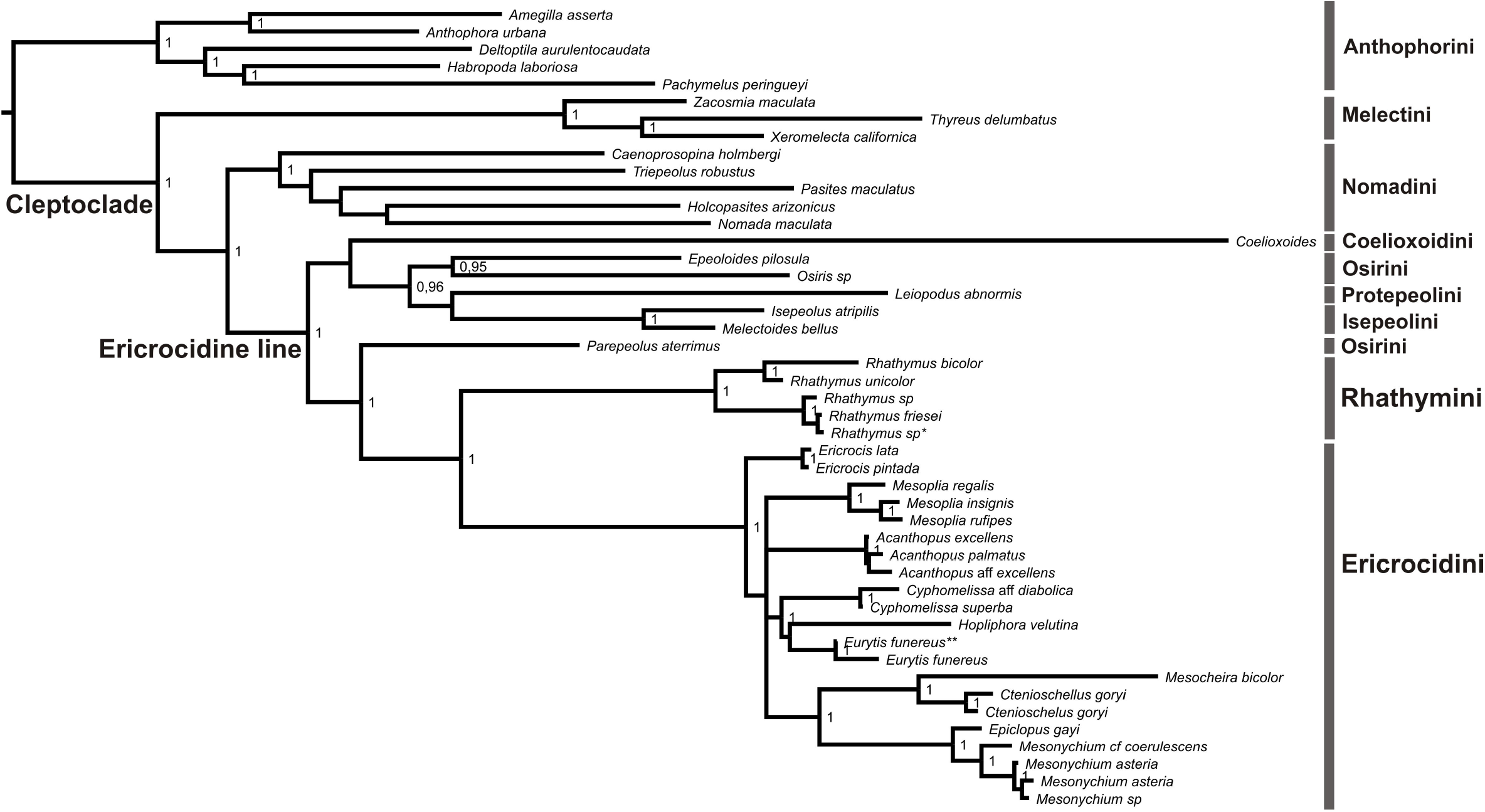
Bayesian phylogenetic tree based on the combined analysis of four nuclear markers and one mitochondrial(46 taxa and 4419 aligned nucleotides) for Ericrocidini plus Rhathymini and other tribes of the cleptoclade (*sensu* Cardinal et al.(2010)) rooted on Anthophorini. Bayesian posterior probabilities (>95) are shown at nodes. In GenBank the following species are identified as *\*Nanorhathymus* sp *\*\*Hopliphora velutina*

The bee tribe Ericrocidini is highly supported as monophyletic (Fig. 3; 1.00 Bayesian posterior probability [BPP], 100% bootstrap support [BS]), as well as its sister relationship to Rhathymini (1.00 BPP; 100% BS), which is also strongly supported as monophyletic (1.00 BPP; 100% BS). The relationships of the well-established clade Ericrocidini+Rhathymini found here are congruent with other higher-level analyses of Apinae (Cardinal et al., 2010). The clade Ericrocini+Rhathymini in our analysis is sister to *Parepeolus aterrimus* (Osirini) with relatively good support (0.99 BPP; 90% BS) as found in other studies on Apinae (Cardinal et al., 2010; Martins et al., 2014). Other Osirini are grouped with Protepeolini and Isepeolini (Fig. 3). The dubious position of *Parepeolus* makes the tribe Osirini paraphyletic, although without morphological support, as it will be further discussed. Despite some small topological differences, the close relationship found here for Ericrocidini +Rhathymini with Osirini, Protepeolini, Isepeolini and Coelioxoidini, also called the ericrocidine clade (Litman et al., 2013), is consistent with previous phylogenetic molecular based studies. Similarly, we found support for the relationship among the nomadine line (Caenoprosopidini, Epeolini, Ammobatoidini, Ammobatini, Nomadini) (0.96 BPP; 78% BS). In both phylogenetic analyses (ML and BI), Melectini is sister to the remaining cleptoparasitic tribes with good support (1 BPP; 100% BS) as in other analysis (Cardinal & Danforth, 2013), but this position seems rather unstable among apine phylogenies. It appears as sister to Anthophorini in our BEAST analysis (Fig. 4) or as sister to the clade formed by the nomadine and ericrocidine lines in other higher-level apine phylogenies (Cardinal et al., 2010; Martins et al., 2014).

**Figure 4.**
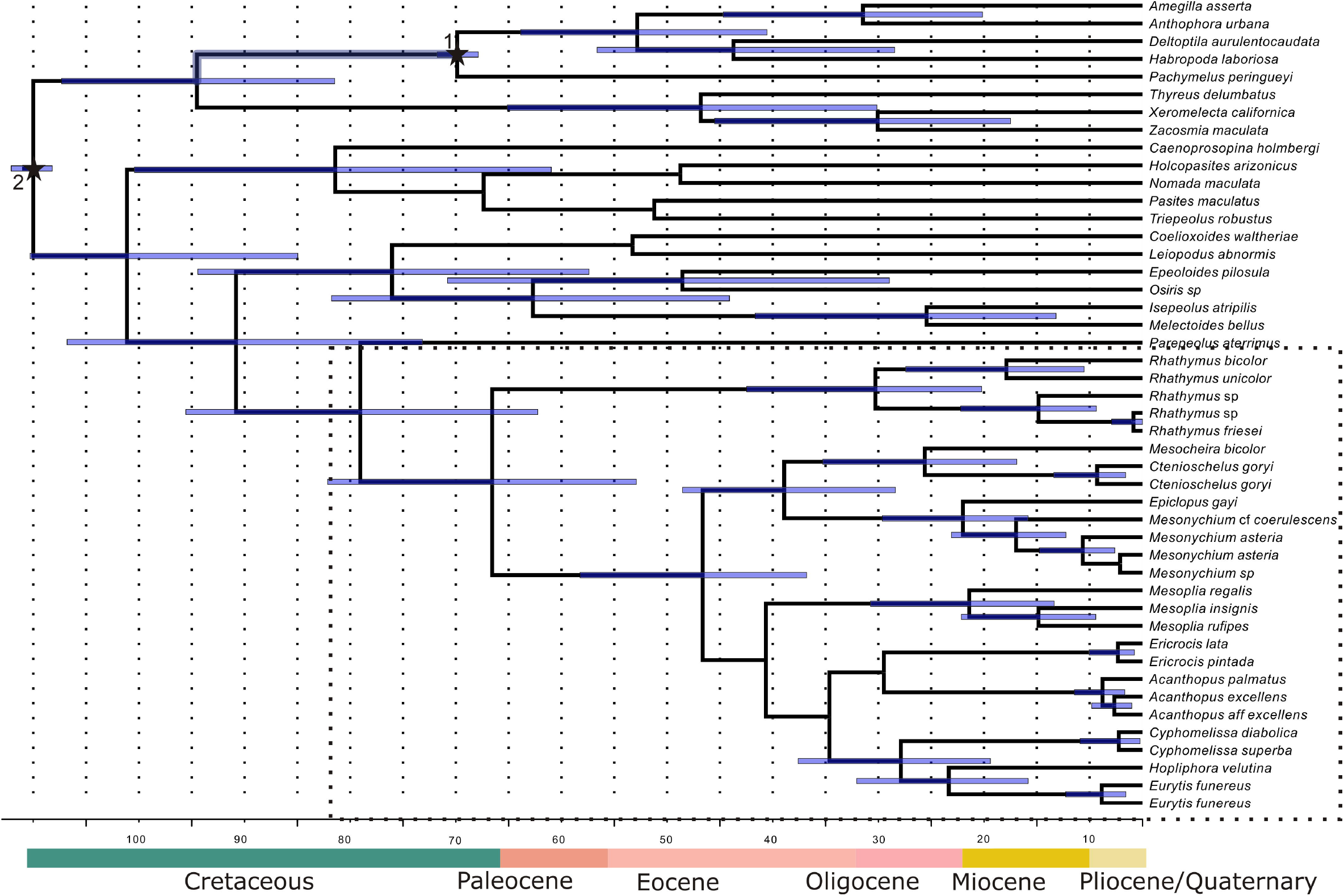
Maximum clade credibility tree derived from BEAST analysis with the same matrix analyzed phylogenetically (Fig. 3 and S1). Node bars represent 95% highest posterior density intervals (HPD) on well-supported nodes. Stars on nodes represent: 1. Fossil calibration point: *Paleohabropoda oudardii* from the Paleocene of Menat, France; 2. Root age constrain: divergence between Anthophorini and the cleptoclade. See material and methods for details on node calibrations. Below the Geological Time Scale (Walker et al., 2012).

As regards the internal relationships in Rhathymini, two main clades were recovered (Fig. 3), both of them with strong support. The first clade contains *R. bicolor* and *R. unicolor* and corresponds to *Rhathymus* s.str., while the second contains the other remaining sampled Rhathymini and corresponds to species that can be placed in *Nanorhathymus* (Engel et al., 2004). *Rhathymus* in its broad sense is composed by twenty described species, widely distributed in the Neotropical region, from Mexico to Paraguay.

All genera of Ericrocidini with ≥2 terminals were supported as monophyletic, corroborating the current genus-level classification (Moure & Melo, 2012a). On the other hand, the basal most relationships in the tribe were not well resolved: in the Bayesian analysis, *Ericrocis* comes out as sister group of the remaining Ericrocidini, with low support, followed by a polytomy with four clades, two being the genera *Acanthopus* and *Mesoplia*, and the other two involving grouping of genera, both of them strongly supported. The group of the four genera *Mesocheira+ Ctenioschelus+ Epiclopus+Mesonychium* represents the first split in Ericrocidini on the ML tree, where it is also strongly supported. *Ericrocis* has a different position on the ML tree, appearing as sister to a clade containing *Cyphomelissa+Hopliphora+Eurytis* and *Acanthopus*, also with low support (Fig. S1).

In contrast to the relatively unstable relationship arrangements at the base of the Ericrocidini, we observe several strongly supported groupings of sister genera, congruent between both trees. The monotypic genus *Mesocheira* and *Ctenioschelus* formed a clade (1.00 BPP; 100 % BS). *Mesocheira bicolor* is widely distributed in Neotropical region (from Central America to Argentina). *Ctenioschelus* is composed by one widely distributed species, *C. goryi*, present in our analysis by two samples, one from Mexico and another from the Brazilian Cerrado, and *C. chalcodes*, occurring in Mexico and Costa Rica. As sister clade to *Ctenioschelus+Mesocheira* we observe another well-supported grouping, composed by *Epiclopus* and *Mesonychium* (1.00 BPP; 100 % BS). *Epiclopus*, which contains four species, is restricted to the Andean region, while *Mesonychium*, with nine described species, is widely distributed in South America. *Eurytis*, which occurs from northern South America to Paraguay, is sister to *Hopliphora*, and both sister to *Cyphomelissa*, all relationships well supported. *Hopliphora, Eurytis* and *Cyphomelissa* have been considered by some authors as a single genus, *Hopliphora* (Snelling & Brooks, 1985; Michener, 2007). *Hopliphora* s.s. possess one widely distributed species (Argentina and Brazil), and *Cyphomelissa* possess four species, two Amazonian (C. *magnifica* and *C. superba*), and two from the Atlantic Forest (C. *commata* and *C. diabolica*).

*Mesoplia* forms a well-supported clade (1.00 BPP; 100% BS) but its sister relationship is not well-defined: in the Bayesian tree it appears as an isolated lineage in the basal polytomy, but in the ML tree it is sister to the clade *Cyphomelissa*+ *Eurytis+Hopliphora* plus *Acanthopus*. All these relationships present low support in the analyses. *Mesoplia* is the most diverse genus in the tribe, with seventeen described species, and presents the widest geographic range, occurring from the southwestern United States (Arizona) to Argentina, including the Greater and Lesser Antilles.

### Divergence times

A molecular clock fossil calibrated tree indicates that the line Ericrocidini+Rhathymini diverged from other cleptoparasitic lineages in the late Cretaceous at 74million years ago (Highest posterior density interval – HPD: 57–90mya) and that they diverged from each other in the Paleocene at 61 mya (48–77) (Fig. 4). The crown age of Rhathymini is estimated as 25 my (15–37) and of Ericrocidini as 41 my (32–53). The ages estimated here for the outgroup sampling, *i.e*. Anthophorini and cleptoparasitic lineages, were very similar to those found by other studies (Cardinal et al., 2010; Martins et al., 2014). All results presented bellow refers to phylogenetically well-supported clades (i.e. BPP ≥0.95; BS ≥70 %) unless stated otherwise.

The two major clades within the Rhathymini have somewhat similar ages, both having differentiated in the Miocene. The age of the clade containing *Rhathymus* s.s. is slightly older (13 my; 5–22) than the other clade (10 my; 4–17). The first split in Ericrocidini, as estimated by the BEAST analysis, separated the lineage composed by the genera *Mesocheira, Ctenioschelus, Epiclopus* and *Mesonychium*, at 34 mya (23–43). *Mesocheira* and *Ctenioschelus* diverged at 20 my (12–30) and *Epiclopus* and *Mesonychium* at 17my (10–24). The crown age for the lineage *Cyphomelissa+Eurytis+Hopliphora* is estimated at 23 (14–32) my, while the split between *Hopliphora* and *Eurytis* is estimated to have occurred at 18 my (11–27). *Cyphomelissa* and *Eurytis* are relatively young, both differentiated in the Pliocene around 3 Mya. The phylogenetic position of the genus *Mesoplia* is uncertain, indicated as related to the clade *Cyphomelissa*+*Eurytis*+*Hopliphora*, although with low support. The BEAST dating analysis placed *Mesoplia* also close to this clade, which in this analysis is sister to *Acanthopus+Ericrocis*. The crown age of *Mesoplia* is estimated as 16 my (8–26), that of *Acanthopus* as 4 my (2–6) and of 2 my (1–5). In any case, considering the many differences between the topology returned by the BEAST analysis and those resulting from the Bayesian and ML analyses, the estimated divergence times within Ericrocidini should be used with caution, in particular for clades not supported in these latter analyses.

## Discussion

### Systematics and divergence times

Our analysis confirms the monophyly of the clade Ericrocidini+Rhathymini plus the sister relationship of this line with *Parepeolus aterrimus* (Osirini) found in previous studies (Cardinal et al., 2010; Martins et al., 2014). We also found support for the ericrocidine line (Litman et al., 2013) as shown in previous analyses. Moreover we show the monophyly of all Ericrocidini genera, corroborating the current taxonomic classification (Moure & Melo, 2012b). In Rhathymini we also found support for recognition of the genus *Nanorhathymus* as proposed by Engel et al. (2004 a, b).

This is the first molecular-based study focused on the tribes Ericrocidini and Rhathymini and we present results that are partly congruent with the previous phylogeny available (Snelling & Brooks, 1985). Snelling & Brooks’ morphological phylogeny, based on a *bauplan* approach, found support for the position of *Ericrocis* as the basal most lineage of Ericrocidini (Snelling & Brooks, 1985), a positioning also recovered here in the Bayesian analysis, although with low support. We also recovered the large clade formed by the genera *Mesocheira+ Ctenioschelus+ Epiclopus+Mesonychium*, but in Snelling & Brooks’ phylogeny *Aglaomelissa* was included and indicated as sister to *Ctenioschelus*. The inclusion of *Aglaomelissa* in our molecular matrix would likely confirm the proximity of this genus to *Ctenioschelus* and *Mesocheira*, considering the strong morphological evidence supporting their close relationship.

Our studies also supported the monophyly of the species treated by Snelling & Brooks as the single genus *Hopliphora* and here represented as the separate genera *Cyphomelissa, Eurytis* and *Hopliphora* s.s. The many morphological differences between these three genera (see key to genera of Ericrocidini in Silveira et al. (2002)) is here reflected in the deep divergence time estimated for them (Fig. 4), equivalent in age to well-established genera in other clades of Ericrocidini.

For both *Acanthopus* and *Mesoplia*, our analyses were inconclusive. The resolution obtained in the ML analysis derives from low-supported branches and therefore seems unreliable as indicator of the relationships of the involved clades. Indeed, the short branches at the base of Ericrocidini suggest a rapid diversification of the main lineages, reflecting a problem that likely will not be promptly solved simply with additional data. Inclusion of additional terminals in future studies should also have a small effect, although more representatives of *Mesoplia*, focusing on a broader representation of this diverse genus, might help in resolving the basal relationships within the tribe.

*Parepeolus aterrimus* was recovered here as sister to Ericrocidini+Rhathymini as found previously and this result implies in a paraphyletic Osirini. Although there is no molecular phylogenetic treatment of the tribe Osirini as a whole, the morphology-based phylogeny provided by Roig-Alsina (1989) places *Parepeolus* within this tribe, as sister to the Chilean genus *Ecclitodes*. The synapomorphies for Osirini are the cervical ventral sclerite and a ventral carina on the forecoxae, while the sister-group relationship between *Parepeolus* and *Ecclitodes* is supported by the enlarged, flattened dorsal branch of the male gonostylus and a bifid ventral branch (Roig-Alsina 1989).

The monophyly of the ericrocidine line—tribes Ericrocidini, Rhathymini, Osirini, Protepeolini, Isepeolini and Coelioxoidini—is once more corroborated here. A morphological character unique to this group is the obliteration of the epistomal suture below the tentorial pit (Melo, unpubl. data). While this character has been used in the diagnosis of some tribes of this lineage, as for example Osirini (Roig-Alsina 1989) and Rhathymini (Engel et al. 2004a), its condition as a putative synapomorphy for the entire line has not been previously recognized.

Divergence time estimates indicated that the origin of the clade Ericrocidini+Rhathymini took place between the end of the Cretaceous and the Paleocene, with the split between them estimated at 61mya. Rhathymini’s crown is slightly younger than that of the Ericrocidini, but both originated between the Eocene-Oligocene border. Among the several cleptoparasitic lineages in long-tongued bees (which totalize at least nine different origins) the cleptoclade represent the oldest origin of this behaviour (Litman et al., 2013), which means Ericrocidini+Rhathymini is one of the oldest lineages of cleptoparasitic bees. As expected, this lineage arose after the origin of their hosts, *Centris* and *Epicharis. Epicharis* line diverged from Centris+corbiculate bees in the Cretaceous, at circa of 90 mya (Martins et al., 2014; Martins & Melo, 2016), much earlier than the differentiation of the ancestral lineage of their parasites. Rhathymini showed an almost coincidental origin to their hosts crown, 25 my (*Epicharis* crown is estimated to be 31 to 28 my (Martins et al., 2014; Martins & Melo, 2016). An even closer match has been found here for Ericrocidini’s crown age and that of their hosts *Centris (Centris* crown is estimated to be 43–41 my) (Martins et al., 2014; Martins & Melo, 2016).

### Diversity and evolution of cleptoparasitism in the Ericrocidini+Rhathymini clade

Except for Protepeolini and Isepeolini which is known to parasitize open cells (by indirect evidence in the latter case), the remaining tribes of the ericrocidine line parasitize closed cells and this seems to be the ancestral state for this clade (Litman et al., 2013). This means they all have hospicidal larvae with morphological and behavioural adaptations to kill the immature host (Rozen, 1989, 1991). Litman et al. (2013) suggested this strategy as a second phase in the evolution of the cleptoparasitic behaviour, derived from the behaviour of parasitizing open cells, in which the adult female bee kills the host larva or egg before laying her own eggs. In a third phase, the adult female deposits her egg in a nest cell that is still open, in process of provisioning, but the hospicidal larva kills other immature present, as found in the nomadine line. The evolutionary scenario within the ericrocidini line, however, is more complex, with Osirini exhibiting a mixture of the first and second phases of Litman’s et al. scheme. In *Protosiris* (Rozen et al., 2006) and most likely in *Epeoloides* (Straka & Bogush, 2007b), the female cleptoparasite kills the host immature (with the sting in *Protosiris* and apparently by eating in *Epeoloides*). At the same time, the larvae of these two genera have hospicidal morphology (Rozen et al., 2006; Straka & Bogush, 2007b). It is possible that the hospicidal morphology exhibited by the cleptoparasite larva evolved first as a weapon against conspecific competitors in situations of multi-parasitized host cells and later was co-opted to kill the host immature, freeing the female cleptoparasite from this task and therefore diminishing the time spent by her to successfully parasitize the host nest.

The ericrocidine line includes most cleptoparasite lineages that use oil-collecting hosts (except *Ctenoplectrina* that parasitize its sister lineage, the oil bee *Ctenoplectra*), *i.e*. Ericrocidini, Rhathymini, Coelioxoidini (parasitizes *Tetrapedia*), and Osirini (parasitizes Tapinotaspidini, in Apinae, and *Macropis*, in the Melittinae). Parasites using oil-collecting hosts occur only in Apinae and evolved one or two times in the large cleptoclade (Habermannová et al. unpubl.). Using oil-collecting hosts obviously limits the host pool, and requires adaptation of larvae to feed on oil, therefore switching to oil-collecting hosts should be less likely than returning back to non-oil colleting hosts (Habermannová et al. unpubl.).

The diversity found in Rhathymini and Ericrocidini is proportional to the diversity found in their hosts. Rhathymini (20 species) and *Epicharis* (35 species) are much less diverse comparatively to their related groups Ericrocidini (44 species) and *Centris* (230 species). This could be the result of the association parasite-host, but also product of many other factors, such as geographical distribution or other limits to the diversification in these groups. The comparative diversity and association between Ericrocidini and *Centris* evidenced the discrepant classification systems currently adopted for the two lineages. While in Ericrocidini, a system with several separate genera is are in use, in *Centris* we observe a more conservative approach, where all species are grouped in a single genus divided in many subgenera. The pattern of host association between Ericrocidini and *Centris*, as well as their antiquity, reinforces the need of treating *Centris* as many different genera due to the number of species, but mainly due to the significant biological differences among the subgenera regarding parasite association, floral host choices, nesting biology and others (see Martins & Melo, 2016).

Whether the higher diversity in Ericrocidini and *Centris* is derived from higher speciation rates or lower extinction rates should be matter for future investigation. It is indeed clear that both groups occupy a wider range, including forests, open plant formations, desert and semi-desert areas, while Rhathymini and *Epicharis* remained restricted to tropical forests. In *Centris* and *Epicharis* this pattern of distribution is associated to their floral host choice. *Centris* species are associated to a broad range of oil-producing angiosperms, belonging to six families, while *Epicharis* collect oil only on species of Malpighiaceae, which are primarily associated to tropical forests (Martins et al., 2015). Probably, oil host plant distribution influences habitat occupation not only of the floral visitors (*Centris* and *Epicharis*) but also of their specialized parasites, Ericrocidini and Rhathymini.

The pattern of host association in Ericrocidini usually follows comparable body size and geographical distribution. As summarized by Rocha-Filho et al. (2009), the relationships are not species-specific, but in some cases all species of a given genus of Ericrocidini parasitize species of the same subgenus in *Centris*, for example all *Acanthopus* attacks nests of *Centris (Ptilotopus)*. Apparently some Ericrocidini genera broke this rule parasitizing more than one subgenus of *Centris*, for example *Mesocheira, Mesoplia, Epiclopus* (Rocha-Filho et al., 2009 and references therein). However this “host broadening” is only apparent and we can observe a preference among the main clades of *Centris:* Melacentris, Trachina and Centris (Martins & Melo, 2016). *Aglaomelissa*, for example, parasitize different species in different subgenera, but all from the clade Trachina.

Comparative studies of the mode of parasitism and larval morphology between Ericrocidini and Rhathymini have been carried by Rozen (1969, 1991), Rozen et al. (2006) and Straka & Bogush (2007a). Rozen (1969) argued for the clade Ericrocidini+Rhathymini based on a greatly elongate labiomaxillary region shared by the mature larvae, according to him “a specialized character that is unlikely to have arisen twice”. After studying the first-instar larvae, Rozen (1991) retracted from his earlier position of a common ancestor between the two tribes due to the many differences presented by their first-instar larvae. One of the most striking differences is the hypognathous head of the first instar larvae of *Rhathymus* compared to the strongly prognathous larvae in Ericrocidini (Rozen, 1991; Rozen et al., 2006). Straka & Bogush’s (2007a) phylogenetic analyses of the larval characters also did not support a sister group relationship between Rhathymini and Ericrocidini.

Considering the robust molecular evidence for the clade Ericrocidini+Rhathymini, one might conclude that the differences in their first-instar larvae are likely adaptations to the distinct nesting biology exhibited by their host bees. The Melectini, the nomadine line and most tribes of the ericrocidini line, including Ericrocidini, have prognathous larvae, with a long and sclerotized head capsule, bearing large elongate mandibles (Rozen, 1991; Rozen et al., 2006), with the Protepeolini and some genera of the nomadine line having an intermediate morphology. Therefore, the Ericrocidini seem simply to have maintained the plesiomorphic condition evolved in the ancestor of the entire cleptoclade.

On the other hand, it is noteworthy to point out that hypognathous first-instar larvae are found only in the tribes associated with oil-collecting hosts, *i.e*. in Coelioxoidini, Osirini and in Rhathymini. If indeed the host’s provisions might exert selective pressures on the cleptoparasites’ immatures (see also Neff & Simpson, 2017) one would wonder why the Ericrocidini do not also exhibit a similar morphology despite attacking oil-collecting *Centris* hosts. Further investigation into this matter should consider the variation observed within *Centris* regarding use of flower oils as larval food. In addition to *C. (Xerocentris)* and some species of *C. (Penthemisia)* that abandoned oil collecting altogether, Vinson et al. (2006) have shown that species of *C. (Hemisiella)* and *C. (Heterocentris)* differ from other studied subgenera in not adding oil to the larval provisions. The few Ericrocidini whose first-instar larvae have been studied (Rozen, 1969, 1991; Rozen et al., 2011) were all obtained from oil-collecting hosts, but addition of floral oils to the provisions has been attested only for *C. (Centris) flavofasciata* (Vinson et al., 1997). A comparative study over a broader range of Ericrocidini species should improve our understanding of their divergence from its sister tribe, the Rhathymini.

## Concluding remarks

This is the first molecular based broadly sampled phylogeny of the Ericrocidini tribe plus Rhathymini, one of the first lineages of cleptoparasitic bees to evolve. We provide phylogenetic evidences that corroborate the current morphologically based classification of both tribes. Moreover, we confirm the relationships among the main lineages of cleptoparatisites in the Apinae, altogether the most diverse group of cleptoparasitic bees, and the monophyly of the ericrocidine line. In this line, we will find most of cleptoparasites attacking oil bees, mostly parasites of open cells and possessing hypognathous first instar larvae. Whether these larval characteristics are related to the use of this alternative floral resource, the oil, by the host is a matter of further investigation. The use of the floral oil in food provisions, and the effects on the cleptoparasites, is still poorly understood. We also ignore the pattern of host association among most ericrocidine line, hampering further conclusions of the evolution of the cleptoparasitism. The long history of host specialization in Ericrocidini+Rhathymini line, provided by field observations, are reinforced by the time of origin of this lineage, almost coincidental to their hosts, *Centris* and *Epicharis*.

## Acknowledgements

We acknowledge Laurence Packer and Jessica Litman for specimen loans. We also thank A. Aguiar, O. Mielke, M. Casagrande, D. Parizotto, P. Grossi, F. Vivallo, H. Werneck, A. J. Donatti and J. T. Souza for providing specimens. AM was supported by scholarships from CNPq and CAPES; DRL (grant 150252/2017-0) and GARM (grants 304053/2012-0; 309641/2016-0) received financial support from CNPq.

## Supporting information Figures

**figures**

**Figure S1.** Maximum likelihood phylogenetic tree based on the combined analysis offour nuclear markers and one mitochondrial (46 taxa and 4419 aligned nucleotides) for Ericrocidini plus Rhathymini and other tribes of the cleptoclade (*sensu* Cardinal et al.(2010)) rooted on Anthophorini. Maximum likelihood bootstrap values (>70) are shown at nodes

**Tables**

**Table S1.** Newly produced sequences used in this study with author names, collecting data, voucher information, and GenBank accession numbers. Entomological collections acronyms: UNB: University of Brasilia, Brasilia, Brazil; DZUP: Entomological Collection Pe. Jesus Santiago Moure, Federal University of Parana, Curitiba, Brazil;

**Table S2.** GenBank sequences used in this study, with species name, collection data and accession numbers.

**Table S3.** Regions of bee DNA sequenced, number of base pairs and related primers and references

